# Different neuroendocrine cell types in the *pars intercerebralis* of *Periplaneta americana* produce their own specific IGF-related peptides

**DOI:** 10.1101/2022.12.10.519892

**Authors:** Jan A. Veenstra

## Abstract

Of the nine genes of the American cockroach, *Periplaneta americana*, coding for peptides related to insulin and insulin-like growth factor, seven show significant expression in the central nervous system as demonstrated by the polymerase chain reaction on reverse transcribed RNA. *In situ* hybridisation shows that five of those are expressed by cells in the *pars intercerebralis*. Antisera raised to the predicted peptides show that these cells are neuroendocrine in nature and project to the *corpora cardiaca*. Interestingly, there are at least three cell types that each express different genes. This contrasts with *Drosophila* where a single cell type expresses a number of genes expressing several such peptides. Whereas in *Drosophila* the neuroendocrine cells producing insulin-like peptides also express sulfakinins, the arthropod orthologs of gastrin and cholecystokinin, in *Periplaneta* the sulfakinins are produced by different cells. Other neuropeptides known to be produced by the *pars intercerebralis* in *Periplaneta* and other insect species, such as the CRF-like diuretic hormone, neuroparsin, leucokinin or myosuppressin, neither colocalize with an insulin-related peptide. The separate cellular localization of these peptides and the existence of multiple insulin receptors in this species implies a more complex regulation by insulin and IGF-related peptides in cockroaches than in the fruit fly.

## 1. Introduction

Insulin is one of the best known and most studied hormones. Its structure is similar to that of the insulin-like growth factors (IGFs), which are also called somatomedins. IGFs are produced and secreted by the liver when triggered to do so by growth hormone released from the pituitary gland and these are the hormones that actually stimulate growth. Insulin and somatomedins belong to a large family of hormones that also include the vertebrate hormones relaxin and INSL-3 among others. Whereas insulin is a metabolic hormone, the other members of this hormone family stimulate growth and different aspects of development and sexual maturation, either in a very general fashion or concentrated on particular details of these processes (Baker et al., 1993; Sonksen and Sonksen, 2000).

Recent analyses of genome and transcriptome sequences suggest that the last common bilaterian ancestor of deuterostomes and protostomes contained a triplet of insulin-like genes (Fig. 1). Those three genes encoded an arch-relaxin, IGF and dilp7 (*Drosophila* insulin-like peptide 7). These three genes likely originated from a gene triplication of an archtype IGF (Veenstra, 2020, 2021a). The genes coding vertebrate relaxin, arthropod gonadulin and ambulacrarian octinsulin descended from the arch-relaxin, while the dilp7 gene persists in insects and other protostomes as well as ambulacrarians (Veenstra, 2020, 2021a). The ambulacrarian data suggests that other, smaller hormones, like insulin, the insect neuroendocrine insulin-like peptides, and the ambulacrarian gonad stimulating substances may have evolved independently from IGF and hence are not direct orthologs of one another, but rather the products of convergent evolution (Veenstra, 2020, 2021a). The term insulin-like peptide and its acronym ilp does not differentiate between these smaller peptides and the triplet consisting of IGF, gonadulin and the dilp7 orthologs. I have therefore proposed to call these smaller peptides short IGF-related peptides (sirps) to make this distinction (Veenstra, 2023). Although sirps are likely descendants of IGF, they are prone to have different functions in different species. Indeed, the experimental evidence suggests that the insect neuroendocrine sirps are growth hormones rather than hypoglycemic or hypotrehalosemic hormones as their structural similarity to insulin might suggest, while the ambulacrarian gonad stimulating substances induce ovulation (Mita, 2019).

**Figure 1.**
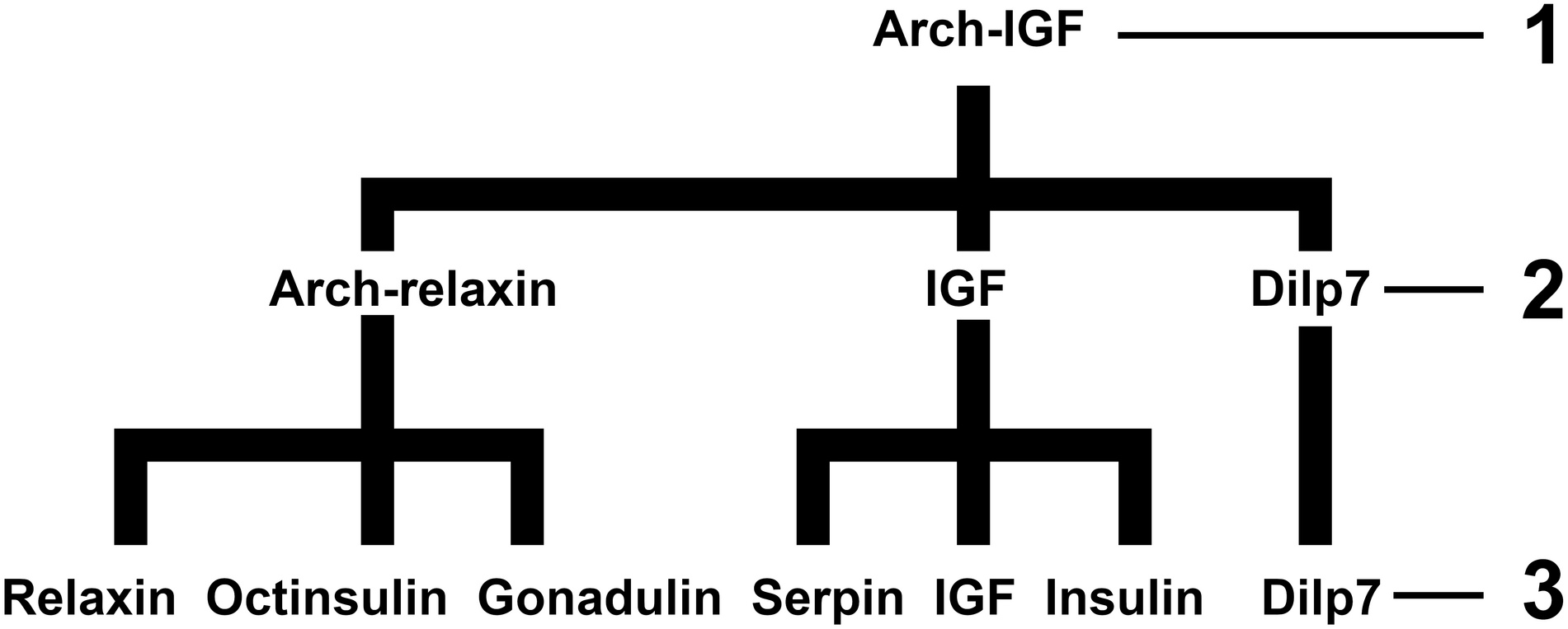
Schematic representation showing how the different extant insulin and IGF-related peptides (3) may be related, as deduced from various protostomian and deuterostomian genome assemblies (Veenstra, 2020, 2021a). The postulated arch-IGF gene from a very early animal (1) underwent a gene triplication, thus yielding three genes coding respectively an arch-relaxin, IGF and a dilp7 ortholog. Those three genes were already present in the last common bilaterian ancestor of protostomes and deuterostomes (2). Ambulacrarian octinsulin, arthropod gonadulin and vertebtate relaxin (as well as several other vertebrate insulin-like peptides) descended from the arch-relaxin gene. The IGF gene persisted and later IGF gene duplications yielded both insect serpins and vertebrate insulin. The dilp7 gene is still present in many protostomians as well as ambulacrarians, but was lost from vertebrates.

Insulin-like peptides are, and have been, extensively studied in the fruit fly *Drosophila melanogaster*. Consequently, we know a lot about their functions in this species which has eight genes coding such peptides that have been called *Drosophila* insulin-like peptides (dilps) 1-8, consisting of five sirps (dilps 1-5), IGF (dilp6), dilp7 and the gonadulin ortholog dilp8 (Brogiolo et al., 2001; Nässel and Vanden Broeck, 2016; Veenstra, 2020). The fruit fly is a highly evolved holometabolous species and its physiology differs significantly from the more basal hemimetabola.

In the latter group the evolution of development and reproduction has led to very different but interesting outcomes, such as viviparous reproduction in the pacific cockroach *Diploptera punctata* and the caste system in termites, which are social cockroaches. It has been noted that termites have three different insulin-IGF receptor tyrosine kinases (RTKs) instead of only one in *Drosophila*. Since activated RTKs play prominent roles in development and growth, it has been suggested that this multiplication of insulin-IGF RTKs might be related to and perhaps have facilitated the development of termite castes (Kremer et al., 2018; Smýkal et al., 2020). As the example of the vertebrate IGF and insulin RTKs suggest, different receptors can induce quite different physiological responses. A recent analysis showed that the American cockroach, *Periplaneta americana*, which is closely related to termites, has even four RTKs (Veenstra, 2020), suggesting that it might be an interesting species to study the various insulin and IGF-related peptides in more detail.

I here report that in *Periplaneta* there are at least three and likely even four different neuroendocrine cell types in the *pars intercerebralis* that each express their own specific sirp genes. This differs significantly from from the fruit fly or the silk worm where there is a single cell type in the *pars intercerebralis* that expresses multiple sirp genes.

## 2. Materials and methods

### 2.1 Antisera

Polyclonal antisera were produced to partial sequences of the different peptides (Fig. 2). This concerns the following peptides: CDGRYNGLPEN (the C-terminal of the B-chain of the predicted *Periplaneta* sirp-1, custom synthesized by Shanghai Royobiotech Co., Ltd [Pudong New Area, Shanghai, China]), pQTRNDRGIVEEC (the N-terminal of the predicted A-chain of the predicted *Periplaneta* sirp-2, custom synthetized by Zhengzhou Phtdpeptides Co., Ltd [Zhengzhou City, China]), CQLPENAERYPFRSRASAVAFP (the C-terminal of the connecting peptide of the predicted *Periplaneta* sirp-3, custom synthesized by Shanghai Royobiotech Co., Ltd), CNGIYNGNPRAPTQ (the C-terminal of the B-chain the predicted *Periplaneta* sirp-4, custom synthesized by Shanghai Royobiotech Co., Ltd), CVLIEEPSDLFKL (the N-terminal of the predicted *Periplaneta sirp-6* precursor related peptide, custom synthesized by Shanghai Royobiotech Co., Ltd), ARSEEDWENAWHRERHTRC (custom synthesized by Pepmic Co., Ltd, Suzhou, China) and CEDWENAWHRERHTGamide (custom synthetized by Zhengzhou Phtdpeptides Co., Ltd). The latter two are part of the B-chain of the *Periplaneta* dilp7 ortholog. Peptides with an N- or C-terminal cysteine were conjugated through this residue using 4-(N-Maleimidomethyl)cyclohexane-1-carboxylic acid 3-sulfo-N-hydroxysuccinimide ester (AK Scientific, Inc., Union City, CA 94587, USA) to bovine serum albumin (BSA). These conjugates were then sent to Pineda Antikörper-Service (Berlin, Germany) who produced polyclonal antisera in rabbits (dilp7 ortholog and sirps −1 and −6), guinea pig (sirp-3) and rat (sirp-4). Antiserum to the predicted *Periplaneta* sirp-2 was produced by Moravian Biotechnology Ltd (Brno, Czech Republic) that also produced the BSA conjugate for this peptide.

**Figure 2.**
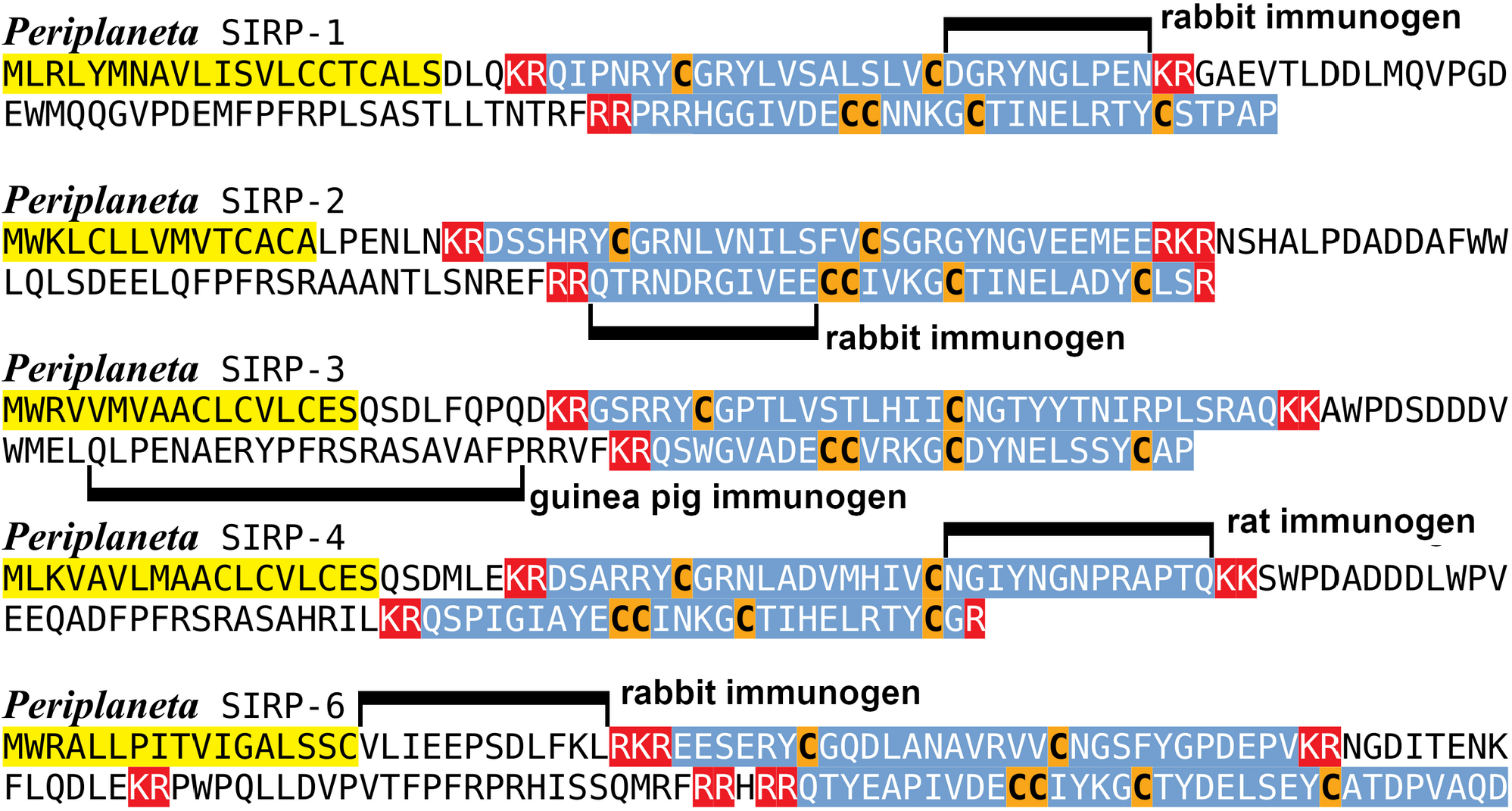
Predicted sequences of *Periplaneta* sirps −1, −2, −3, −4 and −6, indicating their likely processing as well the peptide sequences used for the generation of specific antisera. Yellow indicates the location of signal peptide. Blue highlighting indicates the predicted A and B chains generated after processing of the putative convertase cleavage sites and removal of the dibasic residues, highlighted in red, by carboxypeptidase. The sequences used in antiserum production are indicated by black lines.

IgG from antiserum to sirp-6 were purified using caprylic acid to precipitate serum proteins, followed by ammonium sulfate precipitation. Purified IgG was then labeled with succinimidyl-tetramethylrhodamine and this directly labeled anti-sirp-6 was used in triple labeling experiments.

The following previously described rabbit antisera were also used: *Locusta* neuroparsin (1: 2,000; Bourême et al., 1989), *Periplaneta* sulfakinin (1:20,000; Veenstra et al., 1995), leucomyosuppressin (1:1,000; Schoofs et al., 1993), *Leucophaea* leucokinin IV (1:1,000; Chen et al., 1994), *Manduca* allatotropin (1: 2,000; Veenstra and Hagedorn, 1993), *Drosophila* sNPF-3 (1: 2,000; Johard et al., 2008;) and *Locusta* CRF-like diuretic hormone (1:2,000; Patel et al., 1994).

Secondary antisera: Dylight™488 goat anti-rabbit IgG (minimal cross-reactivity), Alexa Fluoro®488 donkey anti-guinea pig IgG (minimal cross-reactivity), Dylight™549 goat anti-rabbit IgG (minimal cross-reactivity) and Alexa Fluro®647 anti-rabbit Fab fragment were from from Jackson Immunoresearch (Cambridge House, UK).

### 2.2 Affinity columns

Affinity columns were prepared as described previously (Veenstra, 2021b) using PRAESTO^®^ NHS90 Agarose resin (Purolite Ltd, Liantrisant, Wales, UK) and either EEQADFPFRSRASAHRIL (the C-terminal of the connecting peptide of the predicted *Periplaneta* sirp-4) or NGTYYTNIRPLSRAQ (the C-terminal of the B-chain of the predicted *Periplaneta* sirp-3); these peptides were obtained from Zhengzhou Phtdpeptides Co. These columns were used to remove antibodies from the sirp-3 antiserum that might cross-react with sirp-4 and antibodies that might cross-react with sirp-3 from the sirp-4 antiserum.

### 2.3 Insects

*Periplaneta* came for a small colony I maintain on rat chow and water. *Drosophila melanogaster* Canton strain and *D. virilis* were maintained on standard corn meal food.

### 2.4 *Combined* in situ *hybridization and immunohistology*

The combined *in situ* hybridization and immunohistology protocol has been published in detail (Veenstra, 2021b). Briefly, the *in situ* hybridization uses digoxigenin-labeled single strand DNA probes. After fixation in 4% paraformaldehyde buffered in phosphate buffered saline and temporary storage in 70% ethanol at −20 °C, tissues were hybridized with digoxigenin labeled antisense DNA and subsequently exposed to anti-digoxigenin Fab’ fragments conjugated to alkaline phosphatase (Sigma-Aldrich) and peptide primary antisera. After developing the alkaline phosphatase, fluorescently labeled secondary antisera were used to localize neuropeptide immunoreactivity. Pictures of the light microscopy *in situ* hybridization results were inverted, such that the labeled cells become clear and the background black, thus allowing direct comparison with the fluorescent immunoreactivity signals.

### 2.5 SRA analysis

Publicly available short sequence read archives (SRAs) from the American cockroach were downloaded from NCBI (https://www.ncbi.nlm.nih.gov/sra/) and searched for the presence of sequences coding parts of the precursors for the nine insulin-like peptides from *Periplaneta* using the blastn_vdb command from the SRAtoolkit (https://github.com/ncbi/sra-tools/wiki/02.-Installing-SRA-Toolkit) as described previously (Veenstra, 2020). The results are shown in table S1.

### 2.6 RT-PCR

Tissues were dissected under saline and total RNA extracted using Trizol. One μg of total RNA was reverse transcribed in a 20 μl reaction using Moloney murine leukemia virus reverse transcriptase (New England Biolabs, Evry, France) and random primers. One μl of the resulting cDNA was next amplified by PCR for 30 cycles using either OneTaq DNA Polymerase (New England Biolabs) for RT-PCR experiments, or Q5 DNA Polymerase (New England Biolabs) for creating templates for *in situ* hybridization probes. Primers are based on sequences published elsewhere (Zeng et al., 2021), their sequences are listed in Table 1. All primer pairs span introns. Annealing temperatures were those suggested by the New England Biolabs tool (https://tmcalculator.neb.com/#!/main). Sequences of the PCR products were confirmed by automated Sanger sequencing by Microsynth Seqlab GmbH (Göttingen, Germany).

**Table 1.**
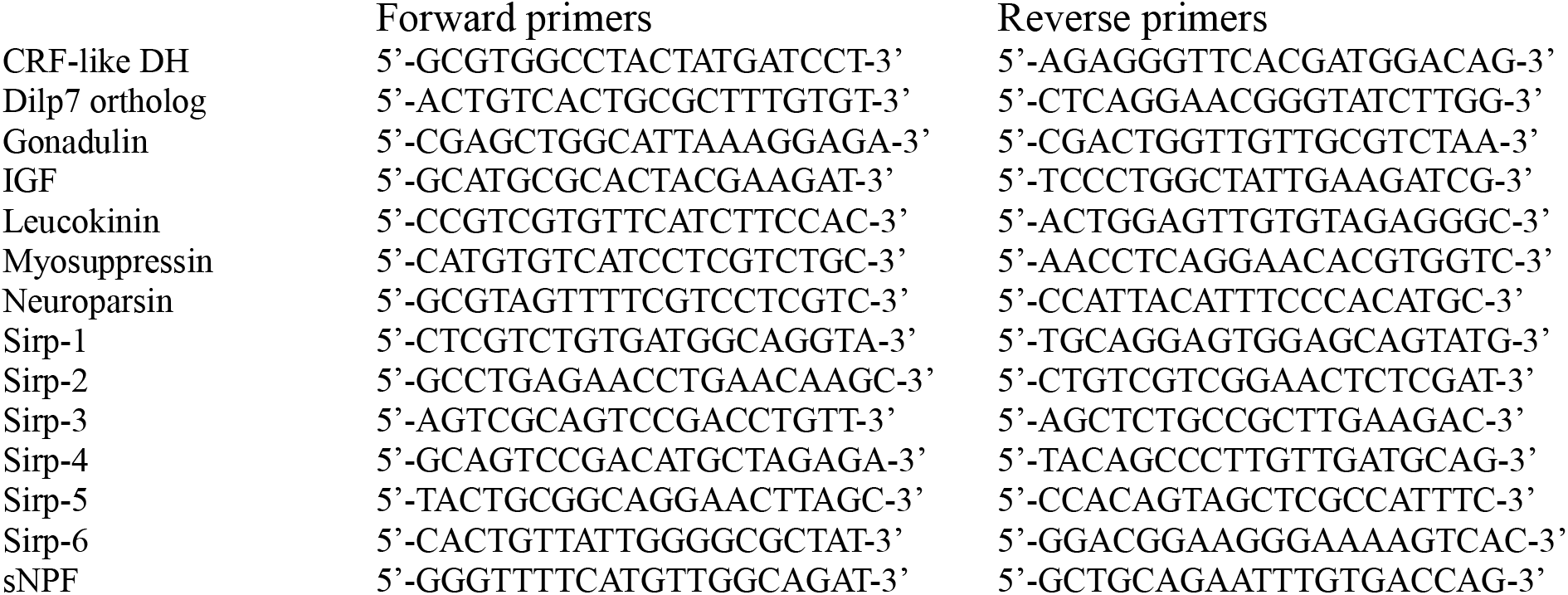
List of primers used for RT-PCR and in *situ* hybridizations.

## 3. Results

Analysis of the brain transcriptome SRAs from the American cockroach (Arvidson et al., 2019) yielded one more sirp than those found in the first genome assembly of *P. americana* (Li et al., 2018), however this gene is present in two more recent genome assemblies from this species (Wang et al., 2022; Zheng et al., 2022).

Analysis of published transcriptome SRAs from *Periplaneta* provided preliminary data on the expression of all nine insulin-like genes (Table S1). The results of the RT-PCR experiments (Fig. 2) concord and included a few additional tissues. Sirps −1, −2, −4 and −6 were only found to be expressed in the brain, while sirp-3 was expressed in the brain as well as in the ovary. Sirp-5 on the other hand was found in all tissues tested, including tissues for which no SRA data is available, like the ovary as well as the collaterial and utricular glands, female and male accessory glands respectively. Gonadulin expression was found in the ovary as well as the testes and utricular glands. Expression in the latter two tissues was variable, in some it was high, in others samples it was absent. A faint band was occasionally also found in the brain. The expression of IGF is strongest in the fat body and ovary, but also significant in the brain, while other tissues reveal very faint bands on the gels (Fig. 3). *In situ* hybridization experiments on sirp-5 in the brain were not successful and neither were such experiments using a gonadulin probe on either the brain, ovary or testes.

**Figure 3.**
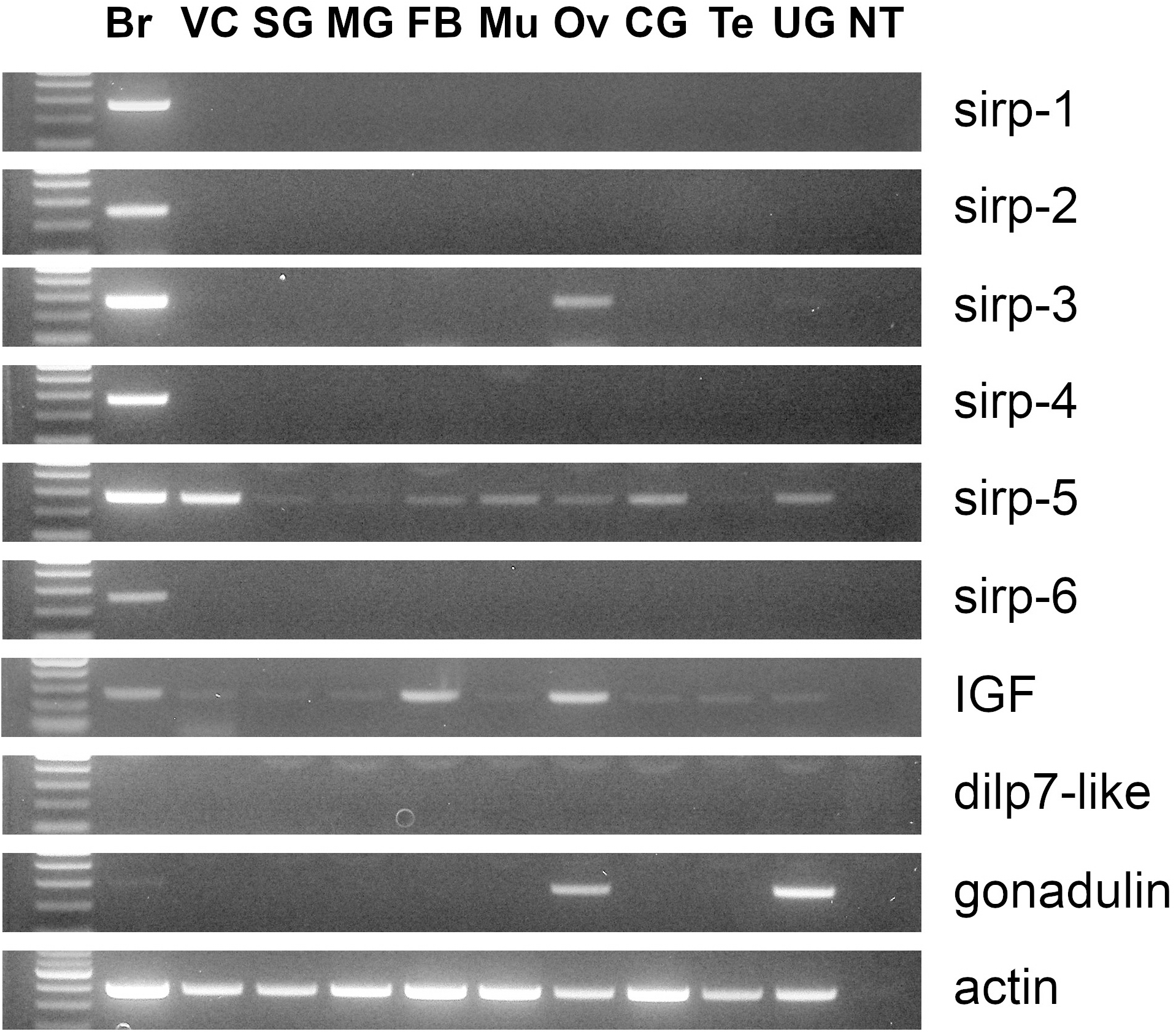
Results of the PCR amplification of cDNA obtained from RNA isolated from ten different tissues: brain, Br; ventral nerve cord, VC; salivary gland, SG; midgut, MG; fat body, FB; thoracic muscle, Mu; ovary, Ov; collaterial glands, CG; testis, Te; utricular glands, UG. The first lane contains molecular weight standards and the last lane a no template PCR control. All primers are intron spanning and no genomic DNA was amplified.

In *Drosophila* the dilp7 ortholog is expressed in a limited number of neurons and it was anticipated that this might also be the case in *Periplaneta*. In initial experiments it was possible to amplify part of the predicted mRNA from the ventral nerve cord and Sanger sequencing results confirmed the amplification product. Nevertheless, *in situ* hybrdization experiments never revealed any positively labeled neurons, while PCR reactions of reverse transcribed RNA from the ventral nerve cord from three further independent experiments failed to yield a positive signal. Two different rabbit antisera to a conserved part of the predicted *Periplaneta* dilp7 ortholog similarly failed to detect any immunoreactive neurons. The primary amino acid sequences of dilp7 orthologs are remarkably well conserved and both antisera recognize dilp7 neurons of both *Drosophila* species tested. Hence, the absence of immunoreactive neurons in the abdominal nerve cord is not due to the antiserum used.

*In situ* hybridization experiments with digoxigenin labeled probes for *Periplaneta* sirps −1, − 2, −3, −4 and −6 all labeled cells in the *pars intercerebralis* suggesting that these might be neuroendocrine cells projecting to the *corpora cardiaca*, like other cells in this brain nucleus. The antisera raised to these peptides showed that they do indeed project there. The *in situ* hybridization signals match the immunoreactivity of the corresponding antisera signals (Fig. 4). Combining the various antisera convincingly showed that sirps −1, −2 and −3 are expressed by different cell types (Fig. 5). *In situ* hybridization signal for sirp-2 colocalizes with both sirp-2 and sirp-6 immunoreactivity showing that these two sirp genes are expressed by the same cells (Fig. 4). Immunoreactivity to sirps −3 and −4 were partially colocalized. The primary amino acid sequences of these peptides are similar, so it seemed possible that the observed colocalization might be due to cross-reactivity of the antisera. These antisera were therefore passed through peptide affinity columns in order to remove cross-reacting antibodies. The antiserum to sirp-3 was passed through an affinity column containing the sirp-4 sequence that is homologous to the antigen used to make sirp-3 antiserum, while the reverse was done for the sirp-4 antiserum. The titer of these preabsorbed antisera was significantly lower than the raw antisera, in particular the sirp-4 antiserum had lost much of its potency and had to be used it high concentrations thereby significantly increasing background staining. Most cells reacting with these preabsorbed antisera were recognized by only one of the two antisera, but there remained a number of cells that were labeled by both antisera. The coding sequences for these two peptides are sufficiently similar (Fig. S1) that cross-hybridization with the respective probes can probably not be excluded. Combining *in situ* labeling for sirp-4 with preaborbed sirp-3 antiserum yields many cells that are identified by only one of the labels, but several cells remain that contain both (Fig. 6).

**Figure 4.**
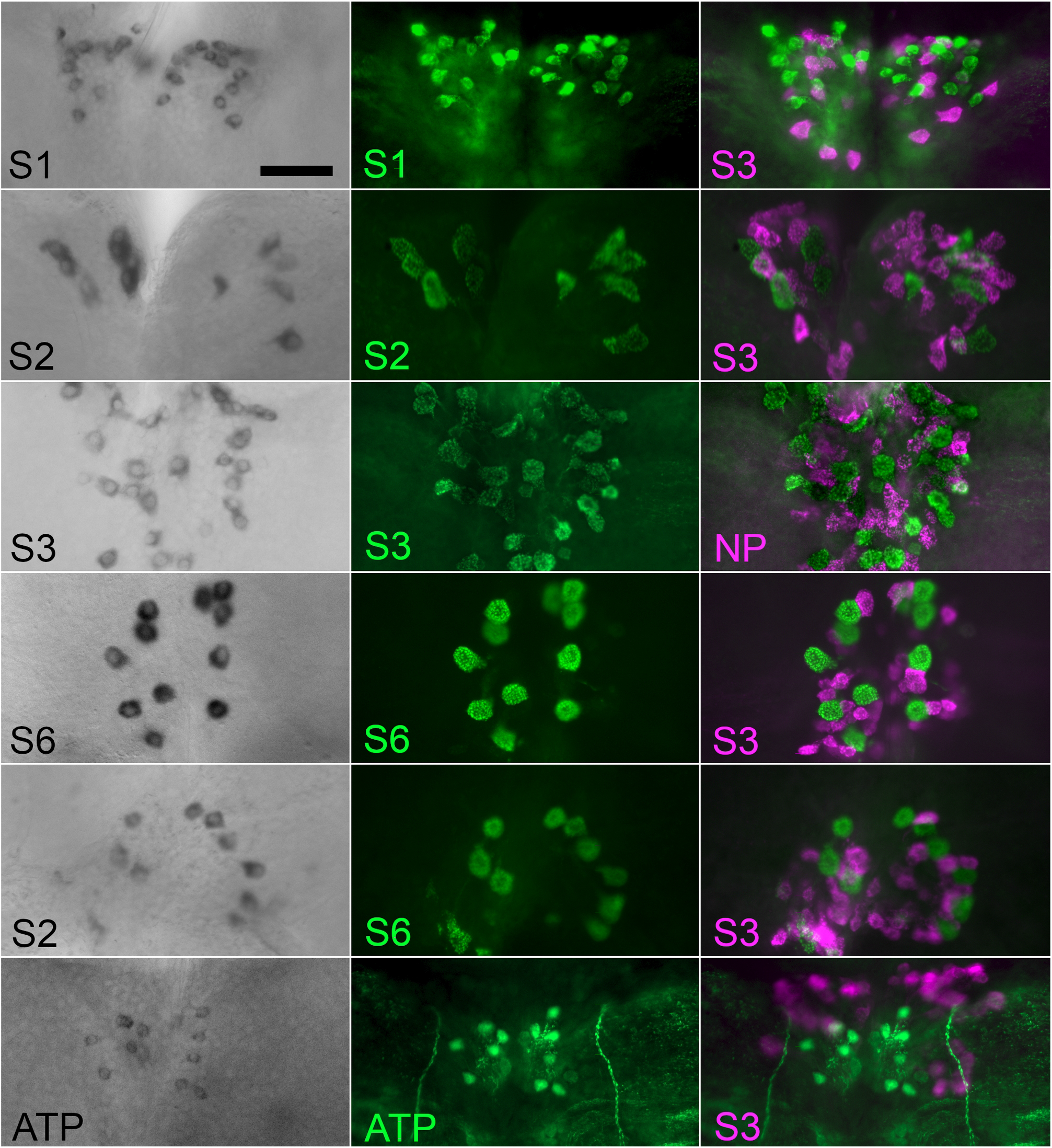
Antiserum specificity. Six whole mounts of *Periplaneta* brains combining *in situ* hybridization with two peptide antisera. The first column shows the results from *in situ* hybridizations of anti-sense probes for sirp-1 (S1), sirp-2 (S2), sirp-3 (S3), sirp-6 (S6) or allatotropin (ATP). The second column shows the immunoreactivity with the corresponding antiserum; all raised in rabbits, except the one against *Periplaneta* sirp-3. The third column shows the labeling with a second peptide antiserum, mostly with guinea pig anti-sirp-3, except one where a rabbit neuroparsin antiserum was used (NP). Note that the *in situ* hybridization signals concord with the antiserum signal for the same peptide, confirming antiserum specificity. Also note that the same cells produce both sirp-2 and sirp-6. Whereas the cells expressing neuroparsin and the various sirps are large, those of the interneurons producing allatotropin are much smaller. False colors were used for the fluorescent signals. Scale bar 100 μm.

**Figure 5.**
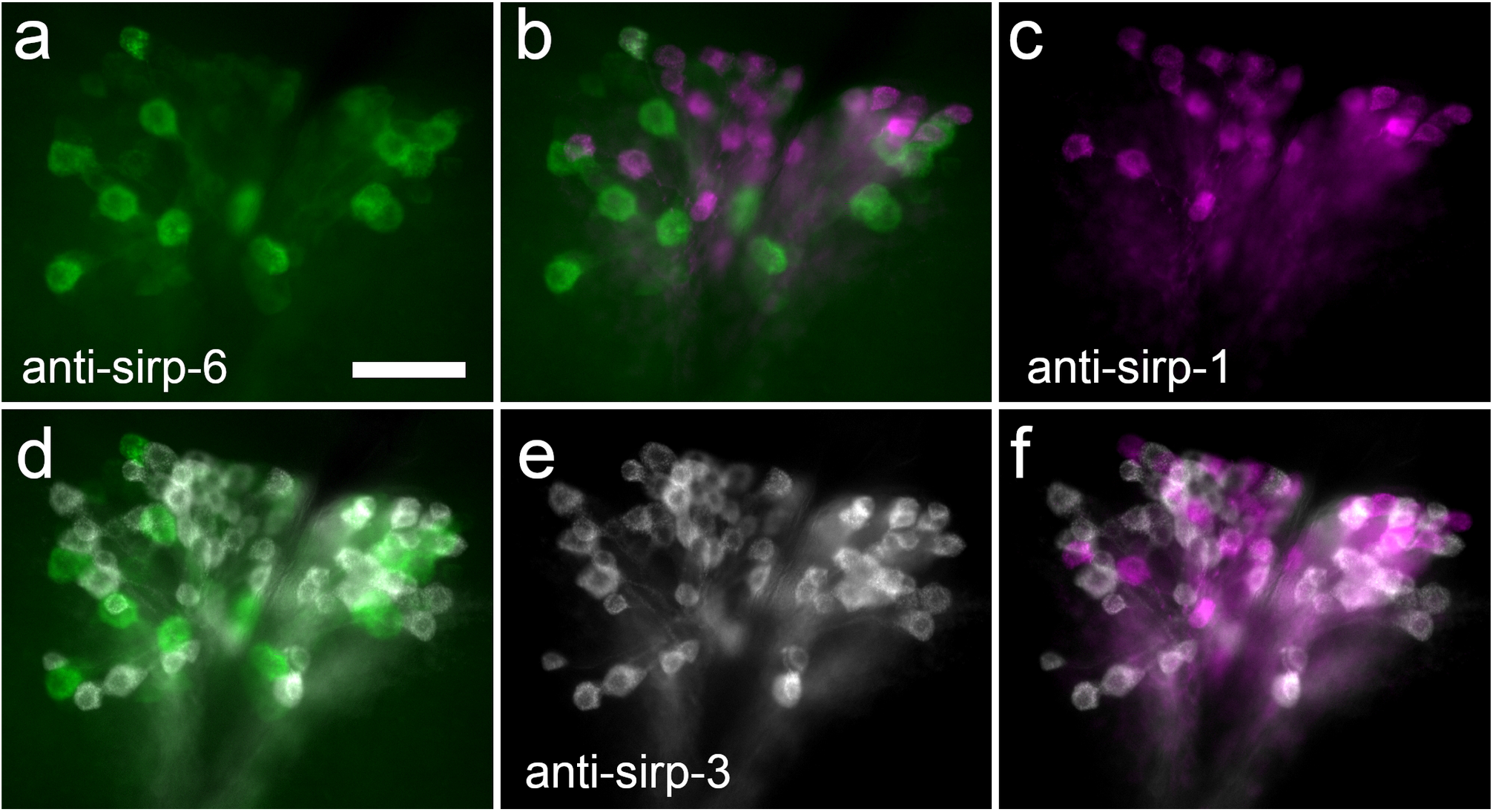
Triple labeling of *Periplaneta* brain with antisera to sirps −1, −3 and −6. Anti-sirp-6 immunoreactivity is green in panels a, b and d, anti-sirp-1 immunoreacativity is magenta in panels b, c and f, while anti-sirp-3 immunoreactivity is white in panels d, e and f. Note that the three antisera recognize different cells. Scale bar 100 μm.

**Figure 6.**
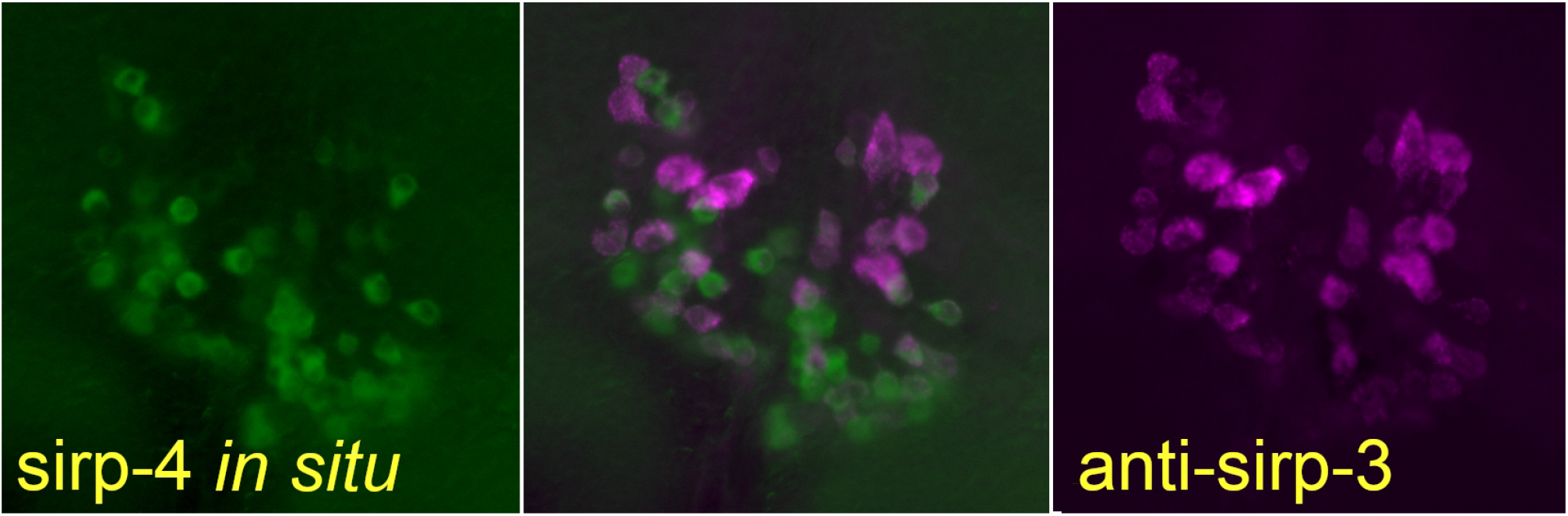
Comparing the sirp-4 *in situ* signal with the sirp-3 immunoreactivity in the *pars intercerebralis*. Note that although the signals largely label different cells, they are not completely distinct. Scale bar 100 μm.

A number of other neuropeptides have been reported to be produced in the *pars intercerebralis*, of this and other insect species, *i.e*. neuroparsin, sulfakinin, myosuppressin, sNPF (short Neuropeptide F), CRF-like diuretic hormone and leucokinin (Tamarelle and Girardie, 1989; Veenstra, 1989; Fusé et al., 1998; Mikani et al., 2015; Kay et al., 1992; Chen et al., 1994). In *Drosophila* the sulfakinin gene is expressed in a subset of the cells expresssing dilps (Söderberg et al., 2012). This raised the possibility that one or more *Periplaneta* sirps might be co-expressed with one of these neuropeptides.

The neuroparsin cells are intermingled with the cells producing the sirps, but are distinct from them. The cells producing the putative diuretic hormones leucokinin and CRF-like diuretic intermingle and are located below the neuroparsin and sirp expressing cells. Those cells are also much smaller than the sirp expressing cells. Perisulfakinin is expressed by about 20 relatively large cells that are often located somewhat at the periphery of the neuroparsin and sirp expressing cells, but these cells do not express any of the sirps. There are 18 myosuppressin expressing cells in the *pars intercerebralis*. This number seems to be fairly constant from one animal to the other. These are commonly located laterally from the other cells and then give the impression of two separate groups, more often they intermingle somewhat with the sirp expressing cells. These cells neither express any of the sirps (Fig. 7).

**Figure 7.**
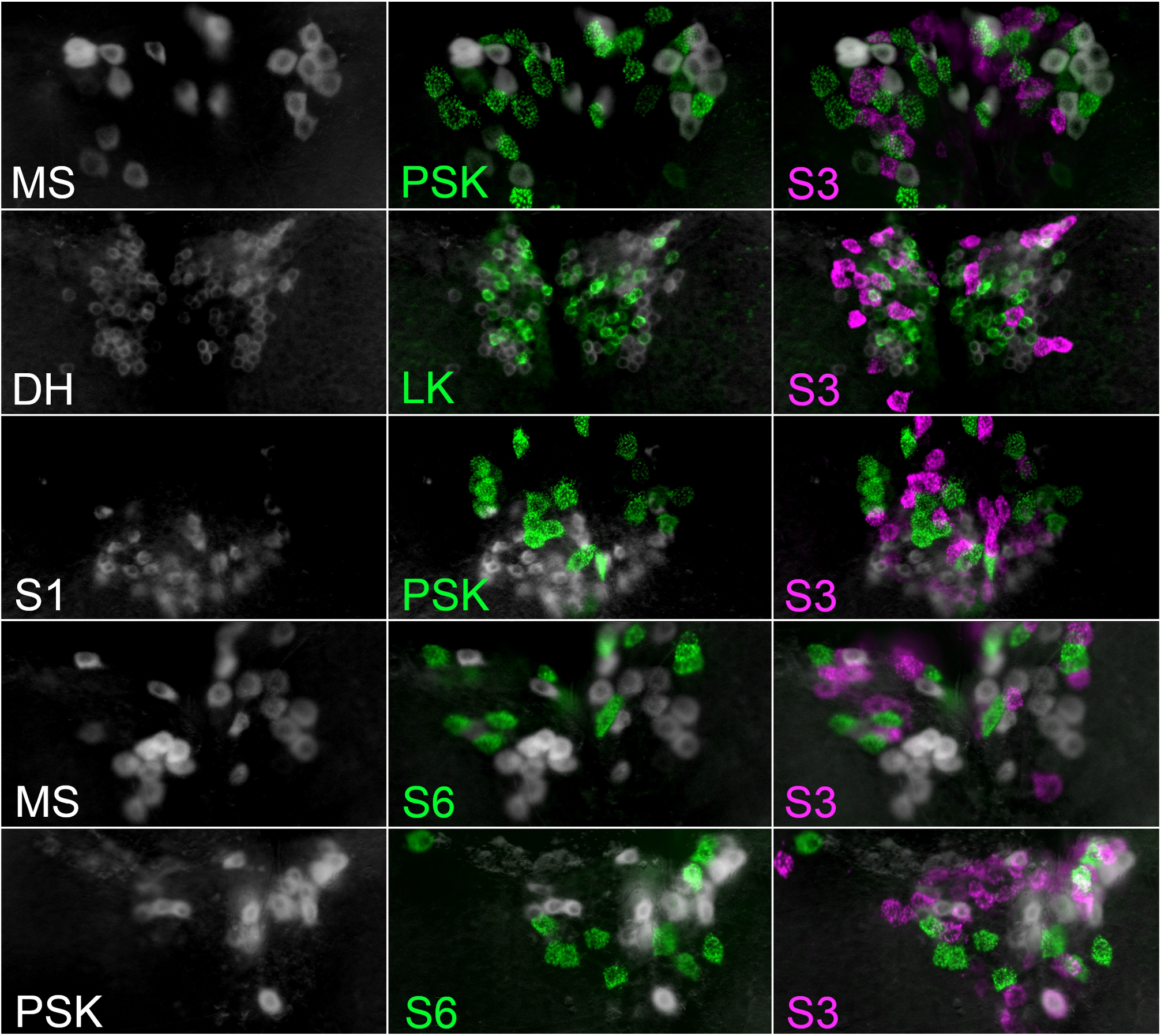
Neuropeptides in the *pars intercerebralis* that are expressed in different cells. Five examples of whole mounts using false colors. The first column shows *in situ* hybridisation, the second column shows the results with the results of a rabbit antiserm added and in the third column the labeling with the guinea pig anti *Periplaneta* sirp-3 is also included. DH, the CRF-like diuretic hormone; MS, myosuppressin; PSK, perisulfakinin; S1, *Periplaneta* sirp-1; S6, *Periplaneta* sirp-6. Scale bar 100 μm.

Others have reported large sNPF-immunoreactive neuroendocrine cells in the *pars intercerebralis* of *Periplaneta* (Mikani et al., 2015), but I was unable to find such cells using antisera to *Drosophila* sNPF-3, an antiserum that is somewhat more specific to sNPF than other antisera to N-terminally extended RFamides. *In situ* hybridization with an sNPF antisense probe neither labeled large cells in the PI, although it does labels smaller sNPF interneurons in the brain. As many antisera to N-terminally extended RFamide cross-react with other N-terminally extended RFamides, it is possible that those large sNPF-immunoreactive neuroendocrine cells express myosuppressin or sulfakinin, that like sNPF also have a C-terminal RFamide. These are however not the only cells expressing neuropeptides in this brain region, small peptidergic interneurons expressing allatotropin can be labeled with both an antisense probe and an allatropin antiserum and it seems likely that there are still other neuropeptides expressed there.

Some of the sirp antisera occasionally labeled cells near the lateral neuroendocrine cells of the brain. In a few preparations the sirp-3 antiserum weakly labeled a neuroendocrine cell en the abdominal ganglia. The location of this cell suggested it might be one of the leucokinin producing neuronendocrine cells, but although the projection of this neuron follows that of the leucokinin cells, it is a different cell. This sirp-3 labeling was inconsistent and these cells were never labeled by the *in situ* hybridization probes

## 4. Discussion

IGF and gonadulin are typically expressed by cells that lack the regulated secretory pathway and are thus expected to be secreted through the constitutive pathway. The results suggest that this is also the case in *Periplaneta*, since these peptides are expressed in tissues – ovary, utricular gland, fat body – that seem to lack cells using the regulated secretory pathway. *Periplaneta* sirp-5 has a similar expression and hence is almost certainly also secreted through the constitutive pathway, whereas sirps −1, −2, −3, −4 and −6 are clearly produced by neuroendocrine cells as shown here.

The presence of four distinct insulin RTKs in the American cockroach *Periplaneta americana* and only one such receptor in *Drosophila* indicated that physiological relevance of insulin-like peptides the cockroach might be significantly different from that in the fruit fly (Veenstra, 2020). As shown here, there are four different neuroendocrine cell types in the brain that all produce their own specific sirps, rather than a single cell type expressing a variety of sirps, as in *Drosophila*, the silk worm or the pea aphid (Brogiolo, et al., 2001; Mizoguchi and Okamoto, 2013; Barberà et al., 2019). This is not the only difference between the fruit fly and the American cockroach. Whereas the expression and physiological relevance of dilp7 specific neurons in the central nervous system is well documented in *Drosophila (e.g*. Miguel-Aliaga et al., 2008; Imambocus, et al., 2021), I was unable to find conclusive evidence for a neuronal expression of its ortholog in *Periplaneta*. Although the question which cells express the *Periplaneta* dilp7 ortholog remains unanswered, its expression seems different from that in the fruit fly.

The big question is of course, why there are so many different sirp expressing cell types and how do their functions differ ? *Periplaneta* is closely related to termites − which are social cockroaches − and similarities between termites and the American cockroach are thus not surprising (Veenstra, 2023). Termites have three different insulin/IGF RTKs and four different sirps, of which one, like *Periplaneta* sirp-5, is expressed in multiple tissues, while the other three are probably expressed by brain neuroendocrine cells. Interestingly, one of latter is also expressed by the ovary (Veenstra, 2023), just like *Periplaneta* sirp-3.

Comparing expression patterns of the four termite sirps it was possible to attribute plausible functions to two of them. The one that is expressed both in the ovary and the brain, called brovirpin in termites, is likely important for vitellogenesis and reproduction in general. One of the other two, called birpin, is probable able to stimulate growth independently from IGF and may be important for the differentiation of larvae into soldiers and reproductives. It is possible that the third termite sirpin would be used by workers to stimulate protein synthesis in the salivary gland, but evidence in support of the latter function is very weak at best. Expression of IGF in termites seems to increase during body growth associated with physogastry that is so prominent in females of the macrotermitidae (Veenstra, 2023).

Given that the expression of *Periplaneta* sirp-3 in both the ovary and the brain resembles that of termite bovirpin it is tempting to speculate that it promotes vitellogenesis and reproduction in general. One or more of the other *Periplaneta* sirps might stimulate growth. It is unknown whether or not in termites the various brain expressed sirps are similarly produced in different cell types nor, if this were the case, whether there are two or three different sirp expressing cell types. Pheromones regulate to a large extent which termite castes will develop. Thus pheromone perception may inhibit neurons, including specific sirp cells. If a termite colony needs neither reproductives nor soldiers, pheromones may inhibit workers to produce the sirps needed to reproduce or develop into soldiers. Such workers could then use available nutritional resources to produce saliva to be fed to other members of the colony by trophallaxis.

As termites are social cockroaches, they were perhaps at least partially prewired to allow their descendants to become social. The closer related to termites, the better that prewiring may have been. *Periplaneta* is very close to cockroaches (Evangelista et al., 2019). In the laboratory cockroaches are kept on protein-rich diets, but like termites they are well adapted to survive on protein-poor diets. It is their *Blattabacterium* symbiont that is able to supply cockroaches essential nutrients like amino acids and vitamins under poor diet conditions (Brooks and Richards, 1955). It is worth noting that when food quality is poor in *Drosophila* larvae, the midgut releases dilp3 as an autocrine hormone to stimulate its growth (O’Brien et al., 2011). Perhaps both cockroaches and termites similarly employ the different sirps to selectively promote some organs/tissues over others, depending on their nutritional conditions. In the case of lack of nutritious food the digestive system might be stimulated, in the case of abundant high quality food, growth or reproduction might be favored. If this were so, the functional differences between sirps in fruit flies and cockroaches might be smaller than they appear at first sight.

## Acknowledgements

I thank Liliane Schoofs, Hans Agricola, Josiane Girardie and Geoffry Coast for generous gifts of antisera many years ago. I am also grateful to Borek Vojtesek, Julio Pineda and their collaborators for making the sirp antisera. The final manuscript benefited from the careful review by three anonymous referees for which I am most grateful.

## Appendix A. Supplementary data

**Figure S1.**
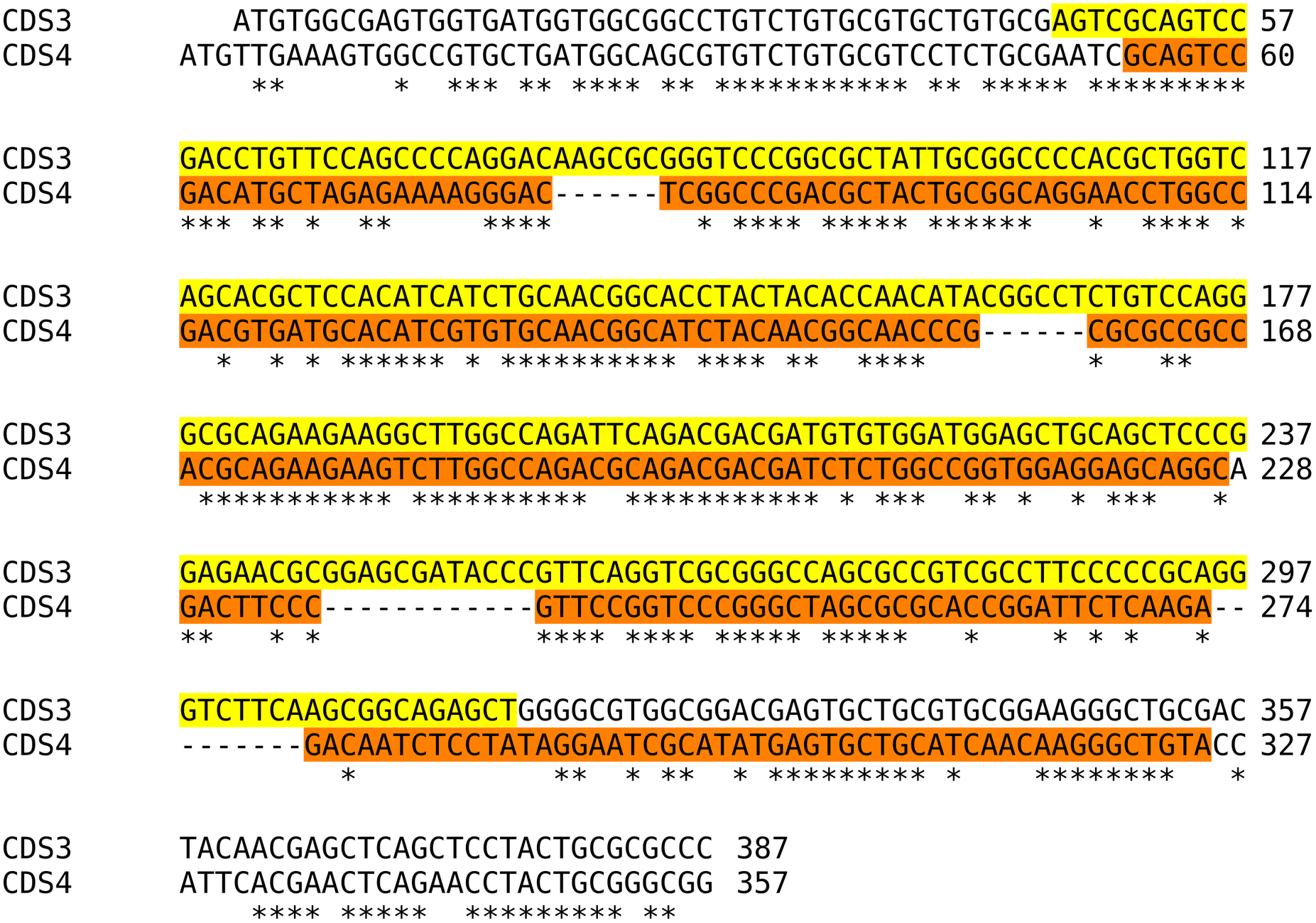
Coding sequences of sirp-3 and sirp-4. The sequences that are amplified and used as templates for the zn *situ* hybridisation probes are colored. Note that these sequences contain very similar sequences.

**Supplementary Table S1.**

| **SRA** | **Sirp-1** | **Sirp-2** | **Sirp-3** | **Sirp-4** | **Sirp-5** | **Sirp-6** | **Gonad.** | **IGF** | **dilp7** |
| --- | --- | --- | --- | --- | --- | --- | --- | --- | --- |
| DRR014884 | 1 | 0 | 4 | 12 | 79 | 0 | 0 | 81 | 1 |
| DRR014885 | 1 | 8 | 29 | 13 | 139 | 0 | 0 | 22 | 0 |
| DRR014886 | 5 | 2 | 16 | 1 | 146 | 0 | 0 | 85 | 2 |
| DRR014887 | 4 | 2 | 10 | 0 | 137 | 0 | 0 | 5 | 0 |
| DRR014888 | 13 | 2 | 34 | 12 | 279 | 0 | 0 | 14 | 0 |
| DRR014889 | 3 | 2 | 11 | 3 | 107 | 0 | 0 | 42 | 0 |
| SRR1184457 | 0 | 1 | 1 | 0 | 557 | 0 | 0 | 3 | 1 |
| SRR1184458 | 0 | 1 | 1 | 0 | 563 | 0 | 0 | 17 | 1 |
| SRR1321695 | 0 | 1 | 0 | 0 | 77 | 0 | 79 | 34 | 0 |
| SRR1322009 | 0 | 2 | 0 | 0 | 157 | 0 | 159 | 57 | 0 |
| SRR2994649 | 21 | 33 | 64 | 38 | 686 | 5 | 0 | 33 | 0 |
| SRR2994650 | 22 | 23 | 319 | 42 | 391 | 4 | 0 | 57 | 0 |
| SRR3056857 | 21 | 33 | 64 | 38 | 677 | 5 | 0 | 33 | 0 |
| SRR3056858 | 22 | 23 | 313 | 39 | 378 | 4 | 0 | 4 | 0 |
| SRR3089536 | 0 | 0 | 4 | 2 | 1499 | 0 | 0 | 31 | 0 |
| SRR3089537 | 0 | 0 | 4 | 0 | 1065 | 0 | 0 | 44 | 0 |
| SRR3089538 | 11 | 15 | 109 | 32 | 1207 | 4 | 0 | 0 | 0 |
| SRR3289663 | 0 | 0 | 12 | 2 | 627 | 0 | 0 | 0 | 0 |
| SRR3289684 | 92 | 0 | 1 | 0 | 79 | 0 | 0 | 2 | 0 |
| SRR3289687 | 6 | 0 | 65 | 0 | 656 | 0 | 0 | 0 | 2 |
| SRR5097509 | 10255 | 11027 | 94318 | 24124 | 14806 | 676 | 0 | 240 | 0 |
| SRR5097510 | 13970 | 10640 | 34115 | 19393 | 24172 | 1026 | 0 | 351 | 0 |
| SRR5097511 | 3174 | 1352 | 4362 | 2406 | 5931 | 180 | 0 | 97 | 0 |
| SRR5097512 | 6 | 76 | 12 | 4 | 23373 | 0 | 0 | 569 | 0 |
| SRR5097513 | 2 | 4 | 11 | 4 | 17436 | 0 | 0 | 0 | 0 |
| SRR5097514 | 2 | 51 | 38 | 12 | 30605 | 2 | 0 | 151 | 0 |
| SRR5097515 | 0 | 0 | 2 | 0 | 20834 | 0 | 0 | 0 | 0 |
| SRR5097516 | 4079 | 3875 | 9774 | 5945 | 4451 | 566 | 0 | 78 | 0 |
| SRR5286150 | 0 | 4 | 66 | 0 | 821 | 3 | 3 | 886 | 4 |
| SRR5286151 | 0 | 0 | 52 | 0 | 1207 | 1 | 13 | 58 | 7 |
| SRR5286152 | 251 | 84 | 342 | 131 | 800 | 8 | 1 | 101 | 6 |
| SRR5286153 | 82 | 34 | 126 | 39 | 1368 | 2 | 0 | 64 | 0 |
| SRR5286154 | 12 | 9 | 47 | 5 | 582 | 0 | 0 | 52 | 0 |

## Notes

### Competing Interest Statement

The authors have declared no competing interest.

### Summary of Updates

Rewritten sections that were unclear, added a figure, changed a figure for clarity.

